# Deep Representation Learning for Temporal Inference in Cancer Omics: A Systematic Review

**DOI:** 10.1101/2025.05.29.656750

**Authors:** Guillermo Prol-Castelo, Davide Cirillo, Alfonso Valencia

## Abstract

Deep learning methods, including deep representation learning (DRL) approaches such as variational autoencoders (VAEs), have been widely applied to cancer omics data to address the high dimensionality of these datasets. Despite remarkable advances, cancer remains a complex and dynamic disease that is challenging to study, and the temporal resolution of cancer progression captured by omics-based studies remains limited. In this systematic literature review, we explore the use of DRL, particularly the VAE, in cancer omics studies for modeling time-related processes, such as tumor progression and evolutionary dynamics. Our work reveals that these methods most commonly support subtyping, diagnosis, and prognosis in this context, but rarely emphasize temporal information. We observed that the scarcity of longitudinal omics data currently limits deeper temporal analyses that could enhance these applications. We propose that applying the VAE as a generative model to study cancer in time, for example, focusing on cancer staging, could lead to meaningful advancements in our understanding of the disease.

**Biographical Note:**

- Guillermo Prol-Castelo is a PhD student at the Barcelona Supercomputing Center and Universitat Pompeu Fabra, where he works on the application of deep learning methods to cancer studies.
- Davide Cirillo is the head of the Machine Learning for Biomedical Research Unit at the Barcelona Supercomputing Center. He is an expert in predictive modeling for Precision Medicine using Network Biology and Machine Learning.
- Alfonso Valencia is the principal investigator of the Computational Biology Group at the Barcelona Supercomputing Center. He is a leading expert in protein coevolution, disease networks and modelling cellular systems.
- The Barcelona Supercomputing Center is a public research center that provides high-performance computing infrastructure to support scientific research in a wide range of fields, including life sciences.

**Key Points:**

- There is a growing interest on the application of deep learning methods, such as Deep Representation Learning (DRL), to cancer studies.
- Cancer is a complex and dynamic disease, whose temporal dynamics are not yet fully captured in omics-based studies.
- mong DRL methods, the Variational Autoencoder (VAE) using omics-based data has been widely used in cancer studies, particularly for subtyping, diagnosis, and prognosis.
- The temporal aspects of cancer progression are often insufficiently captured in omics-based studies, primarily due to the scarcity of longitudinal data.
- Applying the VAE as a generative model to study cancer in time, such as focusing on cancer staging, could lead to significant advancements in our understanding of cancer.

## 1. Introduction

Deep Learning (DL) is capable of capturing complex patterns inherent in health data, including molecular, imaging, and clinical data (1). Given the high-dimensionality of omics data, Deep Representation Learning (DRL) models have emerged as a way to model complex distributions in lower-dimensional representations (2). Moreover, specific DRL methodologies add a generative capability to the model (3). Such is the case of the Variational Autoencoder (VAE), a DRL method that uses Bayesian principles through variational inference to approximate complex relationships between observations and latent variables (4). The VAE, introduced over ten years ago, consists of an encoder and a decoder; the former infers a low-dimensional representation called the latent space, while the latter learns to reconstruct the latent space into the original data. This architecture makes it particularly useful to both model a real data distribution and generate synthetic data (5).

Given the VAE is able to learn non-linear relationships in heterogeneous, high-dimensional data, its use is appropriate for the study of complex diseases. Such is the case of cancer, the second most common cause of death worldwide, with an increasing incidence (6, 7). Well-established techniques for data collection, including omics approaches, have produced large amounts of information, also referred to as big data, about cancer (8). High-throughput molecular and imaging omics data are commonly high-dimensional, noisy, sparse, and heterogeneous. These are also highly informative, including data at different levels of granularity: patient-(9) and/or cell-level (10) resolution; information from different biological sources: genes (11), proteins (12), or metabolites (13); and even spatial information (14). Leveraging DRL methods, such as the VAE, and cancer omics, could have a translational impact by discovering clinically relevant biomarkers (15) and patient subgroups (16).

Navigating such a vast amount of information becomes even more challenging due to the fact that cancer is a dynamic process (17), adding a temporal component. However, two main challenges arise in modeling time-dependent processes in cancer. First, cancer progression is not uniform across individuals (18, 19), meaning that samples collected at the same nominal time points may reflect different progression states across patients. Second, sequencing-based assays are typically destructive, preventing repeated measurements of the same biological sample. As a result, temporally annotated omics datasets generally contain unaligned samples, derived from either multiple individuals or separate samples from the same individual.

Here we present a Systematic Literature Review (SLR) on the use of DRL, paying special attention to the VAE, in cancer studies with omics data for modeling time-related processes, such as tumor progression and evolutionary dynamics. To do so, we consider the following research questions (answered in section 2 Results):

1. How has representation learning, particularly the VAE, been used to study cancer with omics data?
2. How do cancer studies take longitudinal data into account?
3. How is cancer’s temporal dimension studied with deep learning?
4. Which type of omics are most commonly used in time-related cancer studies?
5. What are the key limitations and challenges in the use of DRL for cancer studies with omics data in a temporal context?

The process of the SLR is summarized in Figure 3 (see Methods for details), and our main insights are summarized in Figure 1. We found 440 papers related to our search terms and reviewed their titles and abstracts. These reveal that the most common use of DRL and VAEs in cancer studies are related to subtyping, diagnosis, and prognosis. From the abstracts screening, we selected 44 papers for full-text review, of which 21 were included in our final analysis (summarized in Table 1). We learned that single-cell omics, with a recent surge of spatially-resolved data, is the most common type of data used in time-related cancer studies, albeit in pseudotime, applying the methods included in our search. There is a lack of longitudinal records in the study of cancer in time, and even the temporally annotated datasets available contain unaligned samples. Stages have been considered as a way to provide a time unit for the study of cancer’s advancement, but these usually consider different patients. Overall, we observed a lack of systematic approaches for leveraging temporal omics data to study cancer progression. To address this gap, in the Discussion we outline approaches to address these challenges, such as leveraging the VAE to synthesize temporally aligned instances across cancer stages.

**Figure 1.**
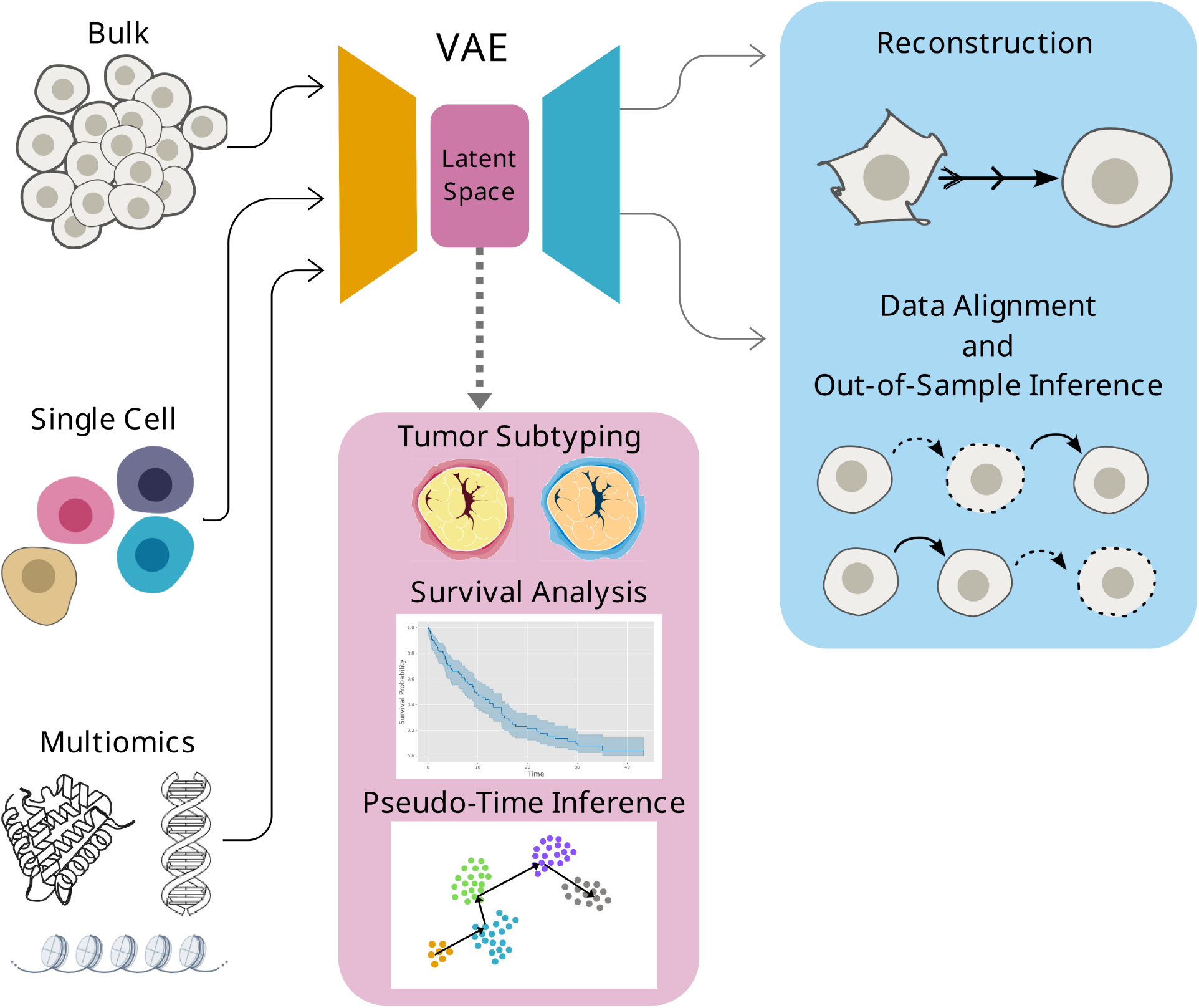
Summary figure of the most common methodologies involving representation learning and cancer studies. On the left side of the figure are shown the most common types of data used to train VAEs particularly—and DRL methods more generally—, in cancer studies. After being passed to the encoder, data is embeded into a lower-dimensional representation, the latent space. The latent space has been commonly used for subtyping, survival anlyses (commonly as part of prognostic analyses), and, when it comes to single-cell data, also for pseudo-time trajectories inference. Possible further applications, shown on the right-hand side of the figure, include leveraging the decoder’s generative capabilities to reconstruct noisy data, and align and infer out-of-sample data, which may be used to study cancer progression in time. However, such applications of the decoder remain underexplored in biomedical research.

**Table 1:**
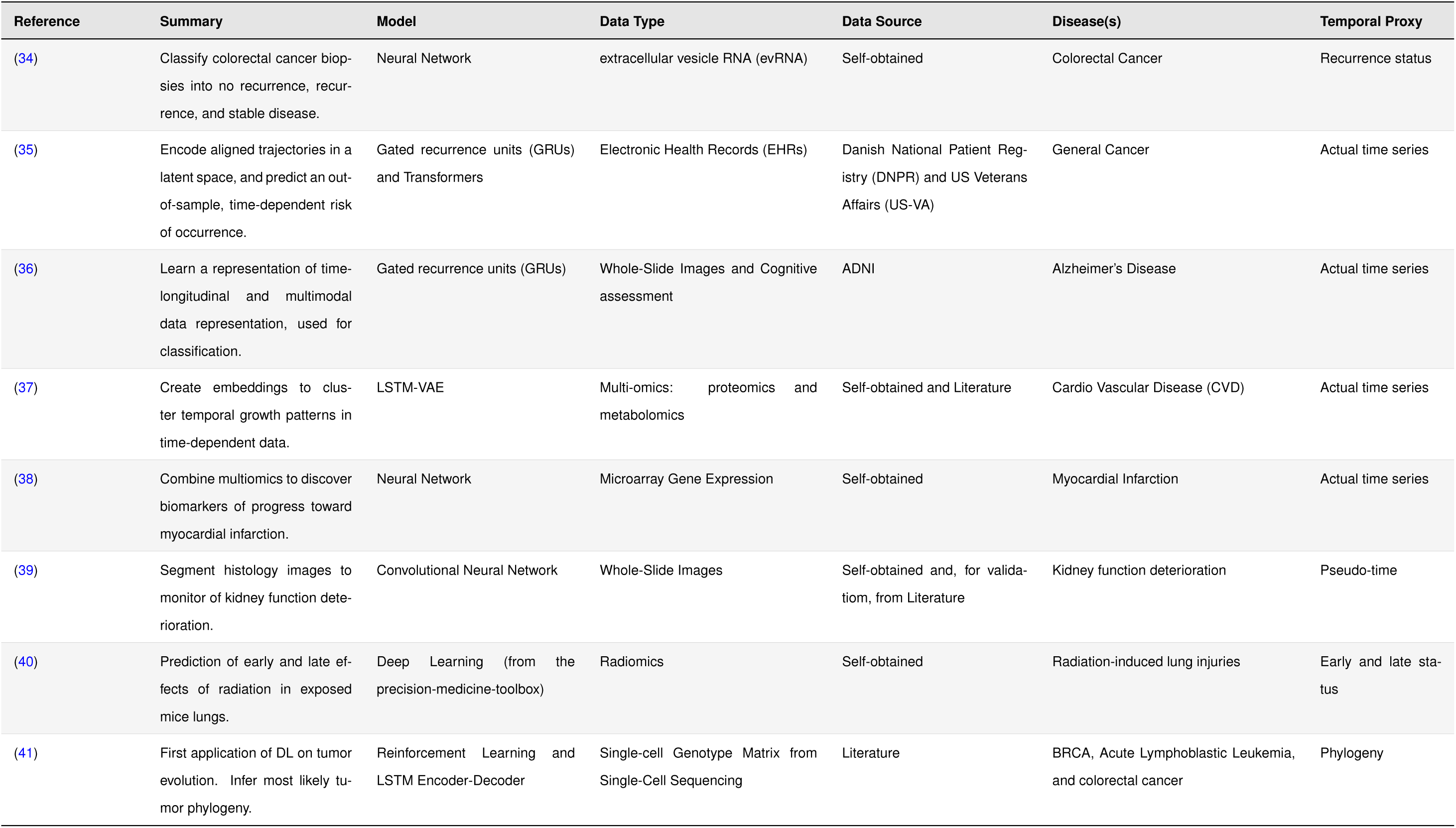
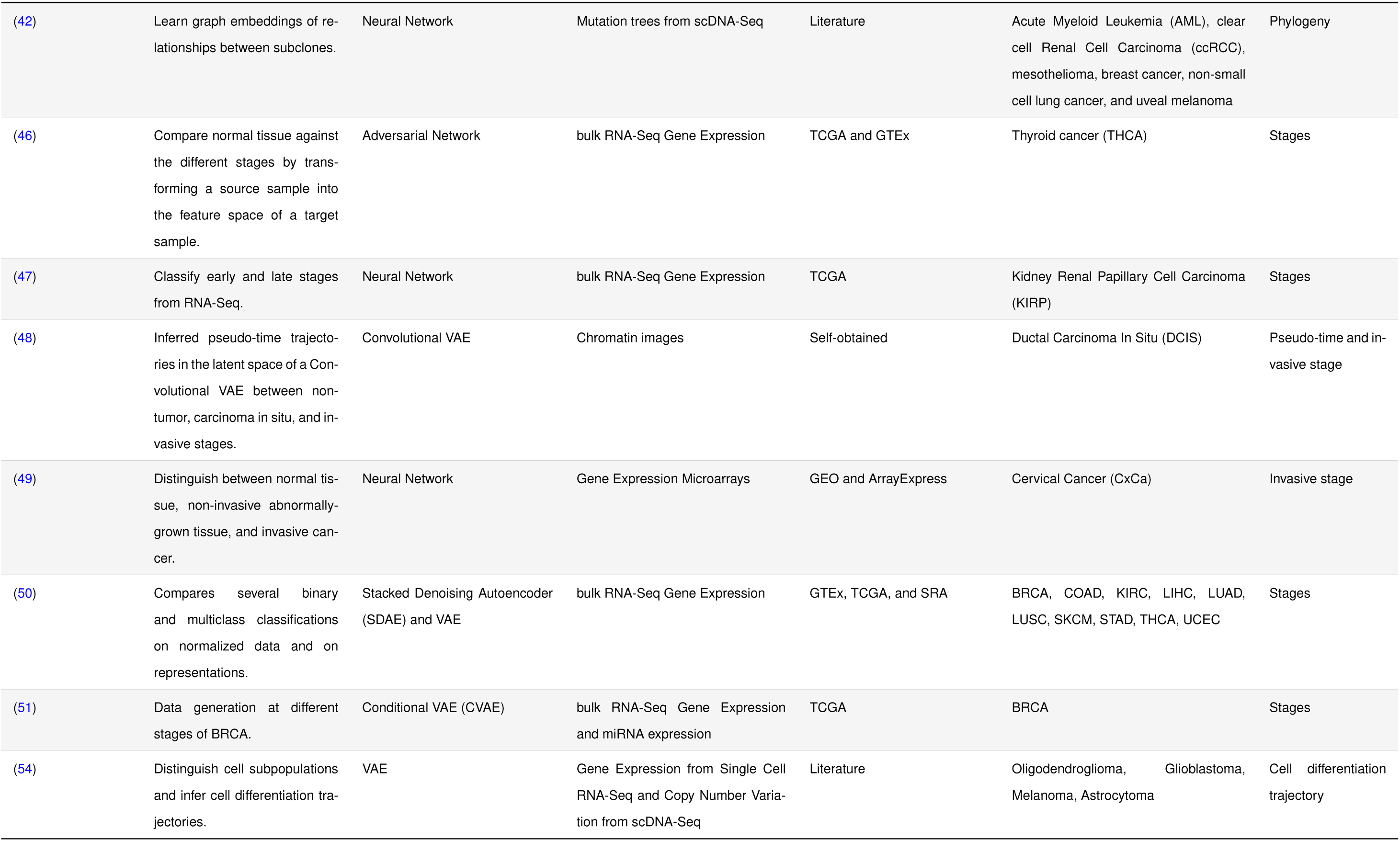
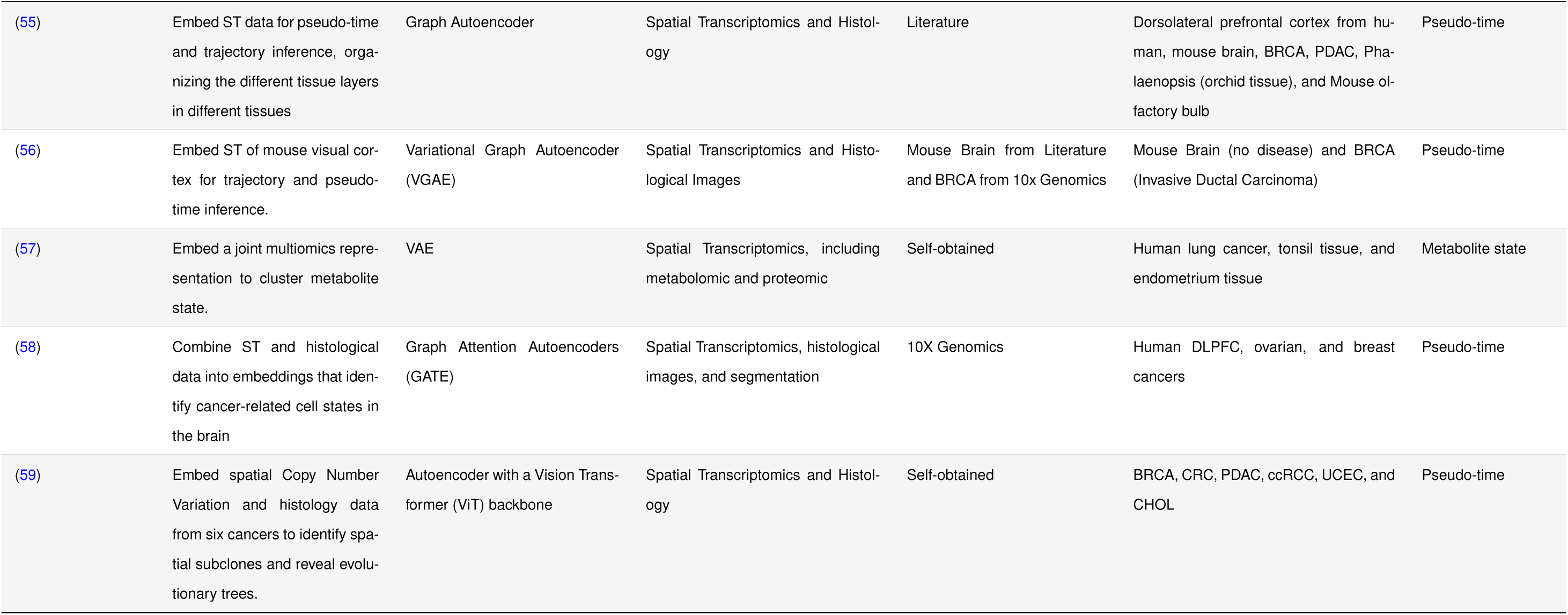
Summary table. This table contains the information of the final list of 21 papers obtained from the Systematic Literature Review (SLR). Papers have been grouped by the used Temporal Proxy, as reflected in the Results section: (34) to (42) provide longitudinal data, (46) to (51) rely on a temporal proxy such as stages, and (54) to (59) use single-cell data and use pseudo-time.

## 2. Results

The five research questions stated on the Introduction are each answered in the following subsections.

### A. VAEs have been used to study cancer diagnosis, prognosis, and subtyping

The most common applications of Deep Representation Learning (DRL) or Deep Learning (DL) in cancer are related to diagnosis, prognosis, or subtyping (Figure 2). Studies of cancer diagnosis aim to distinguish between cancerous and non-cancerous samples (20), or between cancer types (21). Prognosis studies aim to predict the outcome of a disease, which usually involves a survival analysis (22), recurrence prediction (23), or treatment response (24). Subtyping studies aim to identify distinct subtypes, that is, groups into which a given cancer type can be classified (25, 26). All three applications are crucial for developing more personalized treatments that would improve the outcome of the disease.

**Figure 2.**
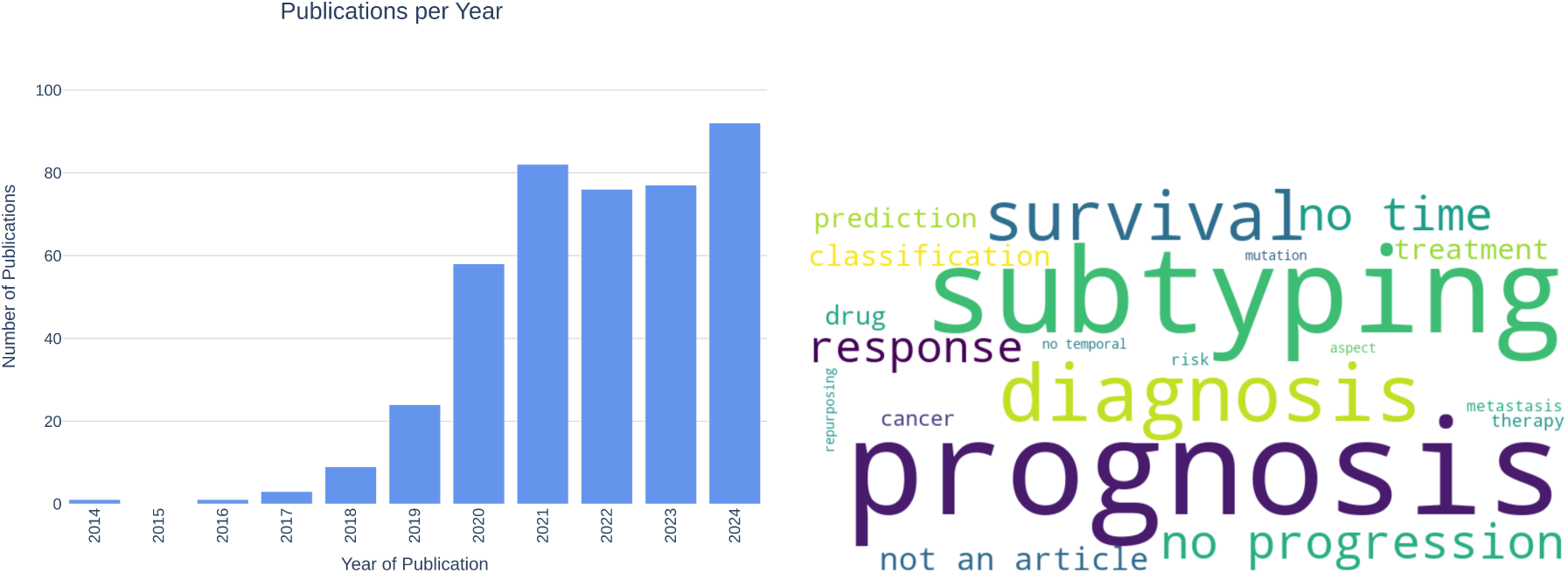
Results from queries. (**A)** Number of publications found, per year. There is a clear increase in relevant publications from 2020. (**B)** Word cloud of exclusion reasons. Most publications use deep and representation learning applied to cancer for subtyping, diagnosis, and studying prognosis and survival.

All of these and other cancer studies rely on the use of high-throughput omics data, which yield information at varying levels of resolution. Bulk omics data have been analyzed using VAEs to learn biologically meaningful low-dimensional representations, enabling the classification of different cancer types in large-scale resources such as The Cancer Genome Atlas (TCGA) (15). Classically, the most accessible data has been bulk data, where the sampled unit is a whole tissue or group of cells. However, the use of single-cell data has been increasing, as it provides more detailed resolution, sampling information from individual cells. For instance, VAEs have been applied to single cell data to detect copy number variations with the objective of probing intratumor heterogeneity (27). Moreover, spatially resolved omics data has recently surged in popularity (28), additionally revealing the spatial distribution of cells in a tissue. Precisely (28) developed an Autoencoder (AE) based methodology capable of distinguishing healthy tissue and different breast cancer types. Omics data also vary in the type of information they provide: transcriptomics data reveal gene expression levels, proteomics data reveal protein levels, and metabolomics data reveal metabolite levels. Besides, omics may be studied in combination, known as multiomics data (29, 22). Namely, (16) used an AE to link multi-omic features to the varying survival outcomes in hepatocellular carcinoma. Recent studies have focused on multiomics and single-cell data, with a recent emphasis on spatially resolved data, given they provide very detailed information and, as such, are expected to reveal new insights into cancer biology. (30) used multi-omics, single-cell data to discern rare cell populations in melanoma. The data included in these studies may go beyond molecular data, including imaging data such as radiomics (31) and histological (32) data. Histological images and DRL have been used to cluster the different areas of lung cancer whole-slide images in order to improve diagnosis (33). Altogether, the surveyed literature shows that DRL methods, especially VAEs, are widely applied to cancer diagnosis, prognosis, and subtyping using omics data, but rarely focus on temporal information.

### B. Longitudinal omics data remain limited in cancer research

The data referenced in the previous section tend to be captured at a single time point. Here, we distinguish between primary and metastatic cancer, which exhibit different growth and behavioral patterns, and focus our analysis on primary tumors. Given the dynamic nature of primary cancer, longitudinal omics data is crucial for understanding disease progression. However, collecting such data from real patients is challenging due to factors including retention difficulties, measurement timing, resource limitations, and ethical considerations. In addition, sequencing-based assays are typically destructive, precluding repeated measurements of the same biological sample. As a result, direct observation of cancer progression in real time is not feasible, either at the patient or cellular level. Even when temporally annotated data are available, inter-patient heterogeneity leads to unaligned observations across individuals. Consistent with these limitations, we observed that the use of longitudinal omics data in primary cancer studies remains scarce.

We found a singular case (34), where extracellular vesicle RNA (evRNA) data from liquid biopsies of colorectal cancer were collected from three cohorts: no recurrence, recurrence, and stable disease. The authors used a DL model to classify temporal subtypes from Gene Set Enrichment Analysis (GSEA) pathways. They were able to find patients that changed subtype during follow-up from the liquid biopsy data, ahead of imaging-based diagnosis. This finding confirms an extra layer of complexity in the study of cancer progression: cancer subtypes may be time-dependent and may be observed through molecular data.

A study following up on cancer patients is (35), where the authors used Electronic Health Record (EHR) data of patients with pancreatic cancer. They used a model based on Gated Recurrence Unit (GRU) and Transformers to embed event features, encode trajectories in a latent space, and predict a time-dependent risk of occurrence. The study shows how representation learning enables out-of sample predictions when longitudinal, aligned data is available.

The use of longitudinal data and its study with DL is more common in other diseases. During our review, we found examples of longitudinal studies on Cardiovascular Disease (CVD), Alzheimer’s Disease (AD), kidney function deterioration, and the effects of radiation. In (36), the authors trained GRUs on radiomics images of the brain and clinical assessments to learn a representation for binary classification of cognitive decline in AD. (37) used a Long Short-Term Memory (LSTM)-VAE trained on multi-omics (proteomics and metabolomics) to embed time-dependent data obtained during cardiac remodeling. The embeddings were then used to cluster temporal growth patterns, determined to be biologically relevant through Kyoto Encyclopedia of Genes and Genomes (KEGG) pathway analysis. Time-dependent microarray data in (38) was collected at different time points pre- and post-myocardial infarction to predict the risk of heart failure. They used a DL model to combine interactomic, molecular, and clinical data, used alongside GSEA to discover biomarkers of progress toward myocardial infarction. Another study, (39), assessed progression of kidney function deterioration with histomorphometry data, applying DL for semantic segmentation.

The authors used trajectory and pseudo-time unsupervised analysis, adapted from single-cell methodologies, to elucidate patterns in disease progression towards kidney failure. Pseudo-time trajectories recapitulated the patterns of damage observed in histology. The authors of (40) collected longitudinal data of blood samples and radiomics from mice lungs exposed to radiation, using DL for survival analyses from the radiomics data. Altogether, these studies highlight the importance of DL and DRL in the study of disease progression, and the potential of longitudinal data to provide insights into the evolution of diseases.

The study of cancer evolution with DL and DRL is common in phylogeny, albeit mostly to discern subclones. A 2020 study (41) claimed to be the first to apply DL to study tumor evolution. Specifically, Reinforcement Learning (RL) was used to infer the most likely tumor phylogeny, and identify tumor subclones. In (42), the authors used unsupervised learning to learn graph embeddings of relationships between subclones. Hence, phylogeny studies of cancer subtypes offer a way to sort subtypes in terms of evolutionary terms. However, while phylogenetic data and studies reflect the evolutionary history of tumors and the subclone ordering, they do not provide explicit temporal resolution in the same way as clinically annotated stages or time-resolved sampling.

### C. Stages as a way to study cancer in a longitudinal manner

Thus far, we have considered actual time as the unit for the study of cancer progression. However, collecting large-scale longitudinal data at regular intervals across multiple patients is often impractical. In addition, tumors evolve at different rates in different patients, so progression is not uniform. In this context, clinically defined stages could serve as a surrogate time unit to study cancer progression. Stages refer to the extent of cancer (43, 44). The most common staging system is the Tumor, Nodes, and Metastasis (TNM), which takes into account the size of the tumor, its spread to nearby lymph nodes, and whether it has metastasized. Thus, there are five stages: stage 0 describes a non-invasive tumor, or carcinoma in situ; stages I through III denote an invasive tumor, the more invasive, the larger the number; while stage IV refers to a metastatic tumor that has spread to distant body parts. Hence, from a conceptual standpoint, cancer stages such as TNM classifications could act as a proxy for temporal progression in the absence of longitudinal omics data, allowing cross-sectional samples to be aligned for trajectory inference, as shown in works such as (45) for GRN reconstruction.

The results from our search highlight a common trend: the difficulty of differentiating cancer stages. As demonstrated in (46), this is addressed by comparing normal tissue to samples from successive cancer stages. Authors used adversarial networks to transform a source sample, normal tissue from TCGA and the Genotype-Tissue Expression (GTEx), into the feature space of a target sample, the different stages, finding several up- and down-regulated genes across stages. Another way to circumvent this issue is to group stages together (47, 48, 49, 50). In (50), data from TCGA was used to classify either stages I and II or stages II and III across different cancers. The classification was compared on real data across different DRL techniques, namely Principal Component Analysis (PCA), Stacked Denoising AE, and VAE. The authors found that representation learning techniques did not improve classification performances. Authors of (47) grouped stages I and II as an “early” stage, and stages III and IV as a “late” stage from papillary renal cancer data. Stages may be simplified even further to distinguish between normal tissue, non-invasive abnormally-grown tissue, and invasive cancer. (49) used DL to distinguish between normal, neoplasm, and cancer samples from cervical cancer microarray data. Besides molecular data, histological images may also be used to classify stages. As shown by (48), a Convolutional VAE was used to learn embeddings of cell states from chromatin images of breast cancer. The authors considered three stages: non-tumor, Ductal Carcinoma In Situ (DCIS), and Invasive Ductal Carcinoma (IDC). The latent embeddings were used to infer a pseudo-time ordering of the stages with Partition-based Graph Abstraction (PAGA), a single-cell trajectory inference method, revealing an ordering from non-tumor to tumor progression. Hence, there is a growing interest in the correct characterization of stages, but its hard separability has led to simplifications in the stages considered.

The lack of separability between stages may be due to the typically unbalanced distribution of samples across stages. In this context, generative modeling may be useful, creating synthetic instances in order to balance out the data. Authors of (51) argued that the use of the Conditional Variational Autoencoder (CVAE) may be used to generate the stages of breast cancer in mRNA and miRNA data from TCGA. They trained the CVAE on the molecular data and used labels of the different stages: I to IV and took solid tissue normal as pseudo-stage 0. The CVAE is then able to generate new, synthetic samples at the trained labels (here, stages). These synthetic stages then showed high separability in a hierarchical clustering analysis. While the study did not report the model loss, it provides an opportunity to reflect on the importance of ensuring that generative models accurately capture the distribution of the original data, and that synthetic samples are representative and properly validated.

### D. Single cell omics data is commonly used to infer pseudo-time trajectories of cancer cells

Single-cell data provide higher-resolution measurements than bulk data, as they profile individual cells rather than aggregate signals from tissue samples. In terms of disease progression, single-cell data can more accurately capture the individual differences between cells within the same tissue. Hence, single-cell data enable analyses that order cells according to transcriptional similarity, allowing the inference of differentiation trajectories. These approaches organize cells along a pseudo-time axis, and a variety of methods have been developed for this purpose, such as Monocle (52) and PAGA (53). During our search, we found that single-cell data was more commonly used for trajectory inference than bulk or imaging data. This data was often processed with DRL techniques to generate lower-dimensional embeddings, which were then used as representations to infer pseudo-time trajectories. For example, the VAE was used by (54) to embed data from Glioblastoma and Oligodendroglioma, and these embeddings were used to infer lineage and differentiation trajectories.

We also found that most of the results from our search that used single-cell data included Spatial Transcriptomics (ST) data, providing information on the spatial localization of cells within the tissue. Besides pseudo-time inference, spatial data also allows to link the different tissue layers. In (55), authors used a Contrast Learning (CL) method to learn latent embeddings of cells and their spots from ST data of the mouse brain. The embeddings were then used for pseudo-time and trajectory inference with PAGA, organizing the different tissue layers. CL was also used by (56) on the mouse visual cortex, combining ST with histological images to learn embeddings of similar spots across tissue regions. PAGA and Monocle were used for pseudo-time trajectory inference, indicating the progression in the developmental process from the white matter layer towards outer layers. (57) used a VAE to embed a joint representation of spatial proteomics and metabolomics data from B cells in the tonsils, and then used these embeddings for metabolite state clustering. The trajectory inference was able to reconstruct the cell differentiation process of B cells in a spatial domain. (58) used a graph attention autoencoder and multiview collaborative learning to embed ST and histological data from human brain slices. The embeddings were used to identify cancer-related cell states: stemness, migration, and metastasis. (59) combined an autoencoder with a visual transformer to obtain embeddings from spatial Copy Number Variation (CNV) and histology data from six cancer types in order to identify spatial subclones and reveal their evolutionary tree. Altogether, the literature indicates the relevance of combining ST data and DRL to relate pseudo-time trajectories with the spatial conformation of healthy and cancerous tissues.

### E. Limitations and challenges

From the studies included in our review, we observed some common trends that limit the study of cancer progression with DRL and DL. First, applications of DL and DRL in cancer studies are mostly centered on subtyping, diagnosis, and prognosis, which typically do not explicitly model the longitudinal aspect of the disease. Studies that do take into account a temporal aspect tend to rely on single-cell data. These approaches typically focus on cellular-level dynamics rather than patient-level progression. Compared to bulk data, single-cell data adds more per-sample noise, as it includes a level of cellular heterogeneity that is not present in bulk data. Moreover, single-cell studies frequently capture relative cellular progression based on transcriptional similarity, rather than the true temporal dynamics of the disease. Finally, clinically defined stages, even when used as proxy time units for cancer progression, are difficult to classify accurately, making robust inference of stage-wise progression from existing omics datasets challenging.

These limitations imply some challenges in the study of cancer progression inference from omics data. First, the lack of longitudinal data is a major challenge, primarily because collecting repeated samples from patients over time is difficult due to logistical, ethical, and clinical constraints, including the impact of ongoing treatments on disease progression. Even when data is collected longitudinally, which is usually more frequent in experimental animal data than human data (60, 61), the rate of cancer progression varies across individuals, so measurements taken at the same nominal time point may not correspond to the same disease stage. A second challenge is the limited availability of methodologies to infer real-time progression. The use of generative models to augment existing data may help address this issue, potentially enabling inference of out-of-sample behavior and effectively introducing a temporal dimension to the dataset. However, this approach leads to a third challenge: the scarcity of data available for model validation. These challenges must be addressed in order to provide a more accurate understanding of cancer progression, and to develop more personalized treatments.

## 3. Discussion

Deep Learning (DL) is a commonly used methodology to learn complex patterns in health data. Deep Representation Learning (DRL) comprises a specific type of DL methods that have the ability to learn a lower-dimensional representation of complex data distributions. This ability has been combined with generative capabilities in the Variational Autoencoder (VAE), a Bayesian-statistics-based representation learning method. Given the complexity of high-throughput data obtained in cancer research, these methods have been applied to study the disease. However, the dynamic nature of cancer implies multiple temporal aspects that remain unexplored.

In this study, we conducted a Systematic Literature Review (SLR) on the use of DRL in cancer studies with omics data for modeling time-related processes. To encompass the complexity of the topic, we created a query that was adapted to three different search engines (see Methods). Our search returned 440 papers, of which 21 were relevant to our investigation (see Tables 1 and 2 for a summary). We summarized these papers and categorized them into 4 main groups. (1) Discarded papers were analyzed to identify the reasons for their exclusion, revealing that the most common applications of DRL and VAEs in cancer are related to subtyping, diagnostics, and prognosis studies. (2) These techniques have been applied in some diseases, taking into consideration a temporal aspect, but their application in longitudinal cancer studies remains limited. (3) Stages may serve as a proxy to align cancer data in a temporal manner. (4) Single-cell omics data is the most frequently used type in cancer studies, with a recent surge in spatially resolved data; however, these studies arrange cells in pseudo-time. Collectively, these findings reveal that the study of cancer progression in time has not yet been adequately explored in omics-based studies.

**Table 2:** Data and Code Summary Table. This table contains the data and code information of the final list of 21 papers obtained from the Systematic Literature Review (SLR). The columns represent: Reference, Dataset size, External Validation, Data Availability, and Code Availability. For External Validation, we take into account only those methods that use tools other than the traditional train-test-validation split.

| Reference | Dataset size | External Validation | Data Availability | Code Availability |
| --- | --- | --- | --- | --- |
| (34) | 155 plasma samples from 71 patients + 70 healthy controls | Consensus Molecular Subtypes (CMS) classifier applied on the TCGA colorectal cancer (n=647). | GSE255775 | Not available |
| (35) | Denmark: 6.2M patients; US-VA: 3M patients | Cross-application (Denmark to US-VA). | Under request to Danish Health Data Authority and VA Boston Healthcare System IRB | Available |
| (36) | Four modalities of data: cognitive performance (n=802), CSF (n=601), MRI (n=865), and demographics (n=865) | Simulation data of 1,000 samples, 500 of which have all data modalities. | Under request to ADNI | Not available |
| (37) | 72 treatment mice and 84 controls | Pathway enrichment to identify which method enriches more significant pathways. | Not available | Not available |
| (38) | 9 premedicated pigs + 3 controls | Pathway enrichment with GSEA. | GSE34569 | Not available |
| (39) | 1,743 WSIs from 1,043 cases | External cohorts (KPMP and HuBMAP). | Raw WSIs available under request to authors; external data from respective consortia | Available |
| (40) | 36 irradiated mice (18 C57BL6, 18 C3H/HeN) + 36 controls | Histology and pathway enrichment. | Not available | Not available |
| (41) | 1,000 simulation samples and 3 real datasets (BRCA 16, Acute Lymphoblastic Leukemia 146 cells, Colorectal cancer 72) | External datasets with known phylogenetic patterns. | Simulation data available in code repository | Available |
| (42) | 16 groups of synthetic trees, 43 tumor mutation trees of 6 cancers, and 123 Acute Myeloid Leukemia trees | Validation against known cancer evolution biology (linear vs branching patterns). | Available in code repository | Available |
| (46) | 432 THCA tumors + 369 normal tissue samples | None reported. | TCGA-THCA | Not available |
| (47) | 259 KIRP samples + 32 normal tissue samples | None reported. | TCGA-KIRP | Available as supplementary material |
| (48) | 560 tissue microarray (TMA) samples from 122 female patients across 3 disease stages (non-tumor, DCIS, IDC) | Pathologist blind-test assignment of tumor grade. | PSI Public Data Repository | Available |

Table 2: **Data and Code Summary Table.**
| Reference | Dataset size | External Validation | Data Availability | Code Availability |
| --- | --- | --- | --- | --- |
| (49) | 202 cancer, 115 cervical intraepithelial neoplasia, and 105 normal samples from 8 datasets | Two datasets used for external validation. | Multiple datasets from GEO (see Table 1 in the publication) | Not available |
| (50) | Almost 40,000 samples | None reported. | recount2 | Available |
| (51) | TCGA-BRCA: 1072 tumor + 104 normal samples. starBase: 863,066 miRNA-mRNA interactions | Pathway enrichment. | TCGA-BRCA and starBase | Available |
| (54) | 6,523 cells across 4 tumors | Separation of malignant and non-malignant cells; differentially expressed genes coincide with literature. | GSE70630, GSE57872, GSE72056, GSE89567 | Not available |
| (55) | Human Dorsolateral prefrontal cortex (12 slices), human ovarian cancer (1 sample, 3,493 spots), human breast cancer (1 sample, 4,727 spots), mouse hypothalamus MERFISH (1,365 cells) | Cross-species and marker genes. | Human DLPFC, Mouse brain, Phalaenopsis, Human breast cancer, Human PDAC (GSE111672), Mouse olfactory bulb, Mouse hypothalamus | Available |
| (56) | Human Dorsolateral prefrontal cortex (12 slices), rest unspecified: mouse hypothalamus MERFISH, mouse visual cortex seqFISH, human breast cancer (10x Visium), mouse olfactory bulb Stereo-seq | Embeddings validated with Monocle3 and PAGA. | Multiple public repositories: spatialLIBD, 10x Genomics, and literature | Available |
| (57) | Human lung cancer (7 patients, 21 regions, 19,507 cells), human tonsil (2 samples, 11 regions, 31,156 cells), human endometrium (5 regions, 8,215 cells) | Multi-tissue application. | IMC and 3D-SMF image data, proteomics, and metabolomics | Available |

Table 2: **Data and Code Summary Table.**
| Reference | Dataset size | External Validation | Data Availability | Code Availability |
| --- | --- | --- | --- | --- |
| (58) | Human Dorsolateral prefrontal cortex (12 slices), human ovarian cancer (for training, 1 sample, 3,493 spots; for validation, 20 samples with 24,489 cells), human breast cancer (for training, 1 sample, 4,727 spots; for validation, 1 sample, 4,081 cells), mouse primary visual cortex (1 sample, 1,365 cells) | Multi-tissue application. | 10x Genomics (ovarian & breast cancer), spatialLIBD (human DLPFC), ClusterMap (mouse visual cortex), GSE176078 (breast cancer scRNA-seq), E-MTAB-8859 (ovarian cancer scRNA-seq), TCGA via Xena | Available |
| (59) | 131 Visium ST sections from 78 cases across 6 cancer types (54 BRCA, 30 CRC, 23 PDAC, 12 RCC, 5 UCEC, 7 CHOL) | Co-detection by indexing (CODEX) samples matching. | HTAN dbGap phs002371.v3.p1, HTAN DCC Portal (WUSTL Atlas) | Available |

The temporal capabilities and the interpretability of the models used in the 21 relevant papers have been summarized and compared in Table 3. Overall, this comparison highlights a trade-off between explicit temporal modeling (e.g., recurrent and attention-based architectures) and latent-space interpretability and generative flexibility, with VAEs uniquely balancing both aspects in omics-based cancer studies. VAEs offer distinct advantages for temporal omics analysis through their latent space representations. Unlike standard neural networks, VAEs learn structured low-dimensional embeddings that preserve data geometry while enabling generative capabilities (51): synthetic data augmentation, missing value imputation, and trajectory interpolation along temporal dimensions.

**Table 3:** Model architectures. Comparison of key characteristics of the models assessed during the SLR process.

| Model | Temporal Modeling | Interpretability | Reference |
| --- | --- | --- | --- |
| VAE (including convolutional, conditional, graph, and attention-based) | Able to embed several disease stages, and to generate data at learned and intermediate stages. | Interpretable latent space. | (48, 50, 51, 54, 55, 56, 57, 58) |
| LSTM and GRUs | Learns long-term relationships. Flexible time intervals. | Requires specific algorithms (85) | (35, 36, 37, 41) |
| Stacked Autoencoders | Learns very complex relationships between variables in data by concatenating several embeddings. Good for feature extraction. | Limited because of the complex relationships between embeddings. | (50) |
| Transformers | Learns long-range relationships. | Requires attention algorithms. | (35, 59) |
| Neural Networks | Mostly used for classification and image segmentation. | Low, but lots of algorithms available. | (34, 38, 39, 40, 42, 46, 47, 49) |
| Reinforcement Learning | For sequential decision-making. | Needs specific algorithms to analyze. | (41) |

The probabilistic latent space enhances interpretability, both by preserving the original data distribution and revealing feature contributions to latent dimensions (62). The VAE may also be further specialized when used in combination with models specific for sequential learning, such as LSTMs (37). These properties make VAEs particularly well suited, compared to non-generative approaches, for reconstructing cancer dynamics from cross-sectional data. However, the VAE still faces some limitations. First, standard VAEs are typically trained in an unsupervised manner with respect to temporal information; conditional variants, such as the CVAE (51), partially address this limitation by incorporating temporal or clinical metadata during training. Second, the highly nonlinear and heterogeneous nature of cancer progression poses challenges for modeling complex temporal distributions within the assumptions of conventional VAEs. During our review, we observed that, to overcome these limitations, the VAE has been used either in combination with other models, such as LSTMs (37), or designed with more complex architectures, such as Stacked Autoencoders (50). Nonetheless, these approaches often come at the expense of reduced interpretability. Finally, VAEs may struggle with out-of-sample generalization, as latent representations learned from limited or homogeneous cohorts may not transfer robustly across patient populations, increasing the risk of generating spurious or biologically implausible trajectories. This issue is analogous to the phenomenon of hallucinations commonly observed in generative AI models (63).

We have attempted to make our search as comprehensive as possible. For this reason, we used multiple search engines and created comprehensive queries, combining deep representation learning terms, omics data types, cancer terminology, and temporal concepts. These queries were designed to cover relevant keywords related to our search questions, while also considering synonyms and alternative terms for the keywords. However, we acknowledge these limitations may have missed relevant VAE-specific temporal studies using alternative terminology or methodological descriptions and did not return papers that could be relevant in the domain. Namely, we identified a number of articles that are not captured by our specific queries but we decided to discuss anyway here due to their relevance. For example, in (64), the authors used a VAE to study the differences across stages of clear renal cell carcinoma using different layers of multiomics data (DNA methylation, gene and protein expression). They revealed genes that were significantly differentially expressed between early (stages I and II) and late stages (stages III and IV). Although not focusing on cancer, another interesting related approach can be found in (62), where they used a VAE to embed bulk RNA-seq data during mouse Central Nervous System (CNS) development. The VAE identified genes with qualitatively different functional profiles and multi-variate trends. The VAE was compared to other embedding methods (PCA, tSNE, UMAP, and PHATE), and was determined to be the most trustworthy at distinguishing genes with known but different anterior–posterior axis association and groups that displayed similar biological enrichment.

The lack of full matches from our queries may reflect substantial challenges and limitations in the field. Research primarily focused on addressing the high dimensionality of omics data through embedding inference, which is sub-sequently used for downstream tasks. These tasks rarely capture temporal cancer dynamics, except in single-cell studies that leverage embeddings to infer pseudo-time trajectories of cells. However, pseudo-temporal ordering does not provide additional longitudinal information, such as clinically diagnosed stages or the actual elapsed time between cellular states. Such longitudinal omics data is difficult to obtain, and when available, its is often collected during treatment and involve destructive sampling. As a result, omics studies of cancer progression frequently rely on data that is not temporally aligned across patients or samples.

The application of the VAE specifically, and DRL more generally, to model temporal progression in cancer omics studies remains underexplored. The field faces significant challenges. On the one hand, some challenges are not specific to time-dependent datasets. Data collected from cancer patients is often heterogeneous, whether it is because of extrinsic reasons, such as different sequencing technologies available and data-processing techniques used, and/or intrinsic reasons, such as the inherent inter-personal heterogeneity of human biology. On the other hand, some challenges are specific to the temporal aspect. These include unaligned data, either because of missing patients follow-ups, or because of sampling at unique time-points or stages of the disease. Furthermore, additional challenges raise when considering multiomics data, due to missing profiles across different omics layers and due to the integration of clinical data (65).

A critical, specific challenge is the need for validation strategies, which would require proper datasets that include the longitudinal dimension. Potential resources include The Cancer Genome Atlas (TCGA), which provides clinical stage annotations, TRACERx, which tracks patients over time with multi-region sampling (66), Patient-Derived Xenograft (PDX) models (67), and pseudo-temporal information derived from large-scale single-cell atlases (68). Validation of synthetic data necessitates ensuring that generated samples meet three key criteria: fidelity, meaning that the original data distributions and biological structure are preserved; utility, ensuring that synthetic data remain informative for downstream tasks such as subtyping and prognosis; and privacy, preventing the re-identification of patients from generated data (69). When detailed, time-annotated datasets are available, they enable benchmarking trained models on their ability to produce temporally aligned and out-of-sample trajectories, as well as to simulate potential interventions, such as treatment responses. This approach has been implemented successfully in single-cell studies for pseudotemporal reconstruction, where generative models are evaluated against known devel-opmental hierarchies (70, 71). Extending similar frameworks to bulk tumor longitudinal data remains an important goal. Addressing these challenges will require open, curated temporal datasets, standardized validation frameworks for synthetic omics data as well as multidisciplinary collaboration. Developing such resources and practices will be critical to advancing VAEs and DRL from exploratory tools toward validated methods for studying temporal cancer progression.

An important goal of the application of AI to cancer research is to have an impact in the clinical oncological setting. Generative AI is no exception. It has shown significant potential at improving cancer diagnostic (72), enabling multimodal data integration (54), and compensating for insufficient data (35). However, there are significant chal-lenges when it comes to the translation of generative AI to the clinic. A key limitation is the lack of interpretability of model outputs (73). Although explainability techniques such as SHAP (74) and LIME (75) exist, our systematic review indicates that deep generative models are frequently applied without accompanying explainable analyses. Besides, Generative AI relies on the characteristics of the data that it learns from, meaning it could replicate or even amplify potential existing biases (76, 77). These challenges raise ethical and regulatory uncertainties (78) that would require rigorous validation to ensure clinical safety and efficacy (79, 80) in order to prevent synthetic biases and hallucinations.

There has been a recent increase in interest in generative models, which can be leveraged to align data, including in the context of temporal analyses. European projects, such as EVENFLOW, are delving into the application of such methodologies to overcome current limitations in cancer studies. Moreover, synthetic data generation methods based on simulation processes, such as agent-based modeling of cellular dynamics (81, 82), are making strides in integrating physics into generative frameworks to produce more realistic and reliable synthetic data. Other initiatives aim to unite experts across diverse disciplines to tackle the challenges of AI in biomedical applications, including generative AI and cancer, as exemplified by the AHEAD project, which brings together specialists in biomedicine, AI, ethics, law, psychology, and related fields. These approaches, together with investments in generating suitable datasets for validation, could pave the way for the development of personalized treatments that account for individual differences in cancer progression.

## 4. Methods

We undertook this systematic literature review in accordance with the Preferred Reporting Items for Systematic Reviews and Meta-Analyses (PRISMA) recommendations (83).

### A. Search process

We performed a systematic literature review of Deep Representation Learning (DRL) methods, with emphasis on VAEs, applied to cancer omics data for modeling time-related processes. Given the complex nature of the topic and the diversity of possible terms related to our research, we considered the following keywords:

- Variational Autoencoder: We considered its abbreviation, VAE, as well. Since the name may not have been explicitly used in the title or abstract, especially during the years after its publication in 2014 (4), we also included the term “representation learning” and “deep learning”.
- omics: We included the main omics data types, such as genomics, transcriptomics, proteomics, and metabolomics, as well as their combination by including multiomics.
- cancer: May also be referred to as tumor.
- time: To include possible references to studies that consider cancer progression through time, such as longitudinal studies, the queries included the keywords progression, evolution, trajectory, time series and longitudinal.

For this review, we used Google Scholar, Scopus, and Web of Science to search the literature, including results from 2014 through 2024.

Given the keywords, we defined the following queries for each search engine:

- Google Scholar: (“deep learning” OR “representation learning” OR “Variational Autoencoder” OR VAE) AND (omics OR genomics OR transcriptomics OR proteomics OR metabolomics OR “multiomics” OR multiomics) AND (“cancer” OR “tumor” OR “tumour”) AND (progression OR evolution OR trajectory OR “time series” OR longitudinal) since:2014 before:2025
- Scopus: TS=(“deep learning” OR “representation learning” OR “Variational Autoencoder” OR VAE) AND TS=(omics OR genomics OR transcriptomics OR proteomics OR metabolomics OR “multi-omics” OR multiomics) AND TS=(“cancer” OR “tumor” OR “tumour”) AND TS=(progression OR evolution OR trajectory OR “time series” OR longitudinal) AND PY=(2014–2024)
- Web of Science: TITLE-ABS-KEY((“deep learning” OR “representation learning” OR “Variational Autoencoder” OR VAE) AND (omics OR genomics OR transcriptomics OR proteomics OR metabolomics OR “multi-omics” OR multiomics) AND (“cancer” OR “tumor” OR “tumour”) AND (progression OR evolution OR trajectory OR “time series” OR longitudinal)) AND PUBYEAR > 2013 AND PUBYEAR < 2025

### B. Selection Criteria

To keep our search within scope and guarantee good quality, we enforced the following inclusion criteria: (1) Peer-review journal articles and conference proceedings; (2) omics data studies; and (3) papers that study cancer evolution or progression using deep learning and/or representational learning. For the same reasons, we also apply these exclusion criteria: (1) Opinions, letters to the editor, reviews, systematic reviews, and gray literature; (2) non-English language works; (3) papers focusing solely on survival and prognostic clinical analyses without providing biological and/or mechanistic insights; (4) papers lacking temporal characterization. The rationale for inclusion and exclusion criteria was motivated by the scope and objectives of this review, which focuses on peer-reviewed research papers the omics-based deep learning and representation learning for studying temporal aspects related to cancer.

When in doubt about borderline cases, we followed a set of decision rules: (1) We included papers that considered the temporal dimension either explicitly, or implicitly through a proxy (such as cancer stages, pseudo-time inference, or spatially-resolved progression patterns); (2) papers focusing solely in prognosis prediction not using omics data were excluded; (3) studies that included only cross-sectional data were excluded unless they incorporated a temporal proxy to study progression (such as cancer stages, pseudo-time inference, or spatially-resolved progression patterns); (4) non-cancer temporal studies were included if all other criteria were met, given the potential translational implications for cancer research. Moreover, (35) was included because it represents a singular showcase of a DRL architecture with generative capabilities (the Transformer) applied to available longitudinal data.

### C. Data extraction

The search on Scopus and Web of Science was performed through the respective websites. Publish or Perish (84) was used to extract the results from Google Scholar. The searches were limited to 200 results for each query. For each query, a table with information about the publication was downloaded from the respective platform. We then identified the duplicated titles and removed them. Next, we read the titles and abstracts, narrowing down our selection. On the remaining entries a full-text screening was performed. The process is summarized in Figure 3. One reviewer (primary researcher) conducted the systematic review process, with results independently verified by the senior author.

**Figure 3.**
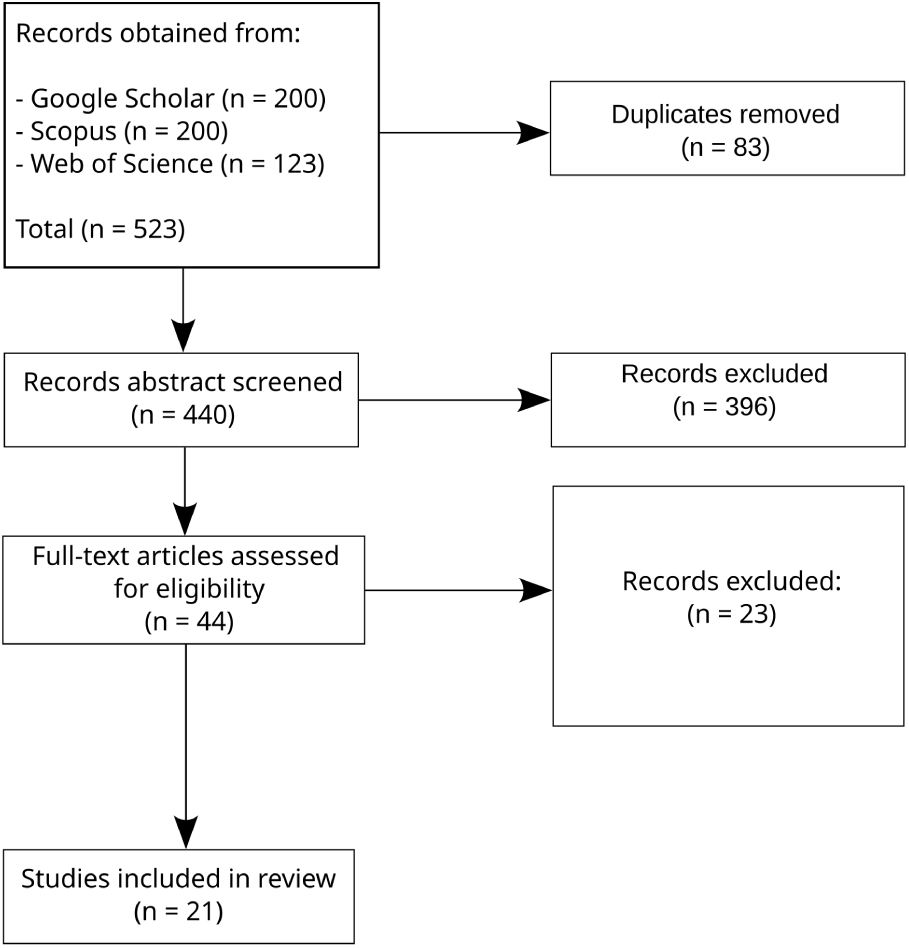
Flow diagram. Summary of the systematic literature review process, following Preferred Reporting Items for Systematic Reviews and Meta-Analyses (PRISMA) guidelines.

### D. Risk of bias

While carrying out this systematic literature review, we identified several possible sources of bias, while taking steps to reduce their impact. The screening process was conducted by one primary reviewer (primary researcher), with a second reviewer (senior author) independently assessing the results to mitigate any subjective bias. While formal inter-rater reliability metrics were not calculated, consensus was reached on all included studies. The data extraction process was designed to mitigate a database coverage bias: the use of three different databases that are not limited to specific fields or literature types (such as pre-print services). We limited the data extraction from Google Scholar to the first 200 results, whereas Scopus and Web of Science returned less than 200 results each from our queries (see Selection Criteria). This is because of the practical feasibility of screening thousands of papers and the diminishing returns of relevance. Even though some relevant papers might have been excluded, the use of three databases was intended to keep the most relevant results. Temporal scope was deliberately restricted to publications from 2014 (corresponding to the introduction of the VAE (4)) through the end of 2024, given our literature search was conducted in January 2025. Publication status bias may also be present: preprints were not excluded, however, only one appeared in our final selection, which may introduce a bias against negative results.

During the screening process, we observed that the majority of journal articles returned by our queries addressed cancer diagnosis, subtyping, and prognosis (see Figure 2), highlighting the considerable research efforts dedicated to these areas. The screening process also revealed a potential application bias within the field, with breast and colorectal cancers being overrepresented (four studies each) relative to other cancer types, and a strong emphasis on analyses using the TCGA dataset.

## 6. Dictionary of Terms

**Table 4:** Dictionary of terms.

| Term | Definition |
| --- | --- |
| **Autoencoder (AE)** | DRL, unsupervised technique that encodes unlabeled data into a lower-dimensional representation, the latent space. It consists of an encoder, which embeds the input data, and a decoder, recreating the input data from the learned embeddings. |
| **Conditional Variational Autoencoder (CVAE)** | A VAE used for semi- or fully-supervised learning. In this case, the encoder and decoder are passed the labels of the input data and learn to encode and decode the input data depending on, i.e., conditioned to, the input data. |
| **Convolutional Variational Autoencoder** | A VAE that entails the use of convolutional layers in the encoder and decoder, making it suitable to learn latent-space embeddings of images. |
| **Contrast Learning** | A supervised DRL method that learns embeddings of data by comparing, or contrasting, between similar and dissimilar instances within the input data. |
| **Deep Learning (DL)** | Discipline within Machine Learning that uses artificial, multilayer neural networks to learn patterns present in a given input dataset. |

Table 4: **Dictionary of terms.** (Continued)
| Term | Definition |
| --- | --- |
| **Deep Representation Learning (DRL)** | Also known as Deep Feature Learning, consists of techniques to automatically estimate a representation of the features included on a given dataset. We use the adjective Deep to remark the fact that we consider Deep-Learnig-based techniques. |
| **Long Short-Term Memory (LSTM)** | Specialized DL model designed to learn when to retain or forget relevant information present in sequential data. |
| **Reinforcement Learning (RL)** | Area of Machine Learning where an automated agent learns a process of decision-making while interacting with an environment and receiving feedback (a reward or a penalty) from an external interpreter. |
| **Variational Autoencoder (VAE)** | DRL technique that follows the AE ability to learn embeddings from unlabeled data, albeit considering a variational Bayesian inference to learn a probabilistic latent space. |

## 7. Data Availability

The results from the queries, as well as the process of literature selection, including independent verification and conflict resolution, are publicly available at the GitHub repository gprolcastelo/SLR-VAE.

## 9. Authors Contribution

G.P.C. developed the systematic literature review, and wrote and revised earlier and consolidated versions of the manuscript. A.V. and D.C. conceived the study and oversaw its execution. Study selection, data extraction, and risk of bias assessment were performed by G.P.C., with results independently verified by D.C. All authors reviewed and contributed to the final manuscript.

## 10. Acknowledgements

This work has been supported by the EU project EVENFLOW under Horizon Europe agreement No. 101070430.

